# A model for estimating traction force magnitude reveals differential regulation of actomyosin activity and matrix adhesion number in response to smooth muscle cell spreading

**DOI:** 10.1101/612267

**Authors:** Sultan Ahmed, Panashe Mabeza, Derek T Warren

## Abstract

Decreased aortic compliance is associated with ageing and vascular disease, including atherosclerosis and hypertension. Ultimately, changes in aortic compliance are driven by altered ECM composition however, recent findings have identified a cellular component to decreased aortic compliance observed in ageing and hypertension. Vascular smooth muscle cells (VSMCs) line the blood vessel wall and VSMC contraction regulates vascular tone and contributes to aortic compliance. Mechanical cues derived from the ECM influence VSMC function, yet whether ECM rigidity influences VSMC force generation remains unclear. In this study, we describe the relationship between VSMC spreading, traction force magnitude and matrix rigidity. Importantly, we show that spreading predicts integrated traction force (integrated-TF) magnitude independently of matrix rigidity. Using linear regression analysis, we have generated a model for calculating integrated-TF from VSMC area. This model closely predicts the integrated traction force measured by live VSMC traction force microscopy. Vinculin staining analysis revealed that spreading strongly correlated with adhesion number per VSMC, suggesting that increased VSMC integrated-TF was due to enhanced matrix anchor points. Further analysis revealed that calculated integrated-TF per adhesion was reduced by matrix rigidity, however, adhesion number/μm^2^ increased, resulting in the average integrated-TF/μm^2^ remaining unaltered. As a result, the integrated-TF/VSMC spreading relationship is independent of matrix rigidity. Therefore, our study has identified and validated a novel model to predict and understand the mechanisms influencing VSMC traction force magnitude.

## Introduction

Decreased vascular compliance is a risk factor in the development of cardiovascular disease, including atherosclerosis and hypertension (1, 2). Rodent models of hypertension display a 4-fold increase in aortic stiffness compared to age matched controls (3). Additionally, the Young's modulus, E, (a measure of material stiffness) of healthy aorta has been determined by atomic force microscopy to be between 10-20kPa, whereas atherosclerotic plaques contain a stiffened fibrous cap (E=60-250kPa) (4). Ultimately, these changes in aortic compliance are driven by altered extracellular matrix (ECM) composition. Elastic ECM components, including elastin, are degraded and non-elastic ECM components, such as collagen-1, accumulate (5, 6). However, recent studies have identified a cellular component to aortic compliance (3, 7–9). Vascular smooth muscle cells (VSMCs) are the predominant cell type in the medial layer of the aortic wall and normally exist in a quiescent, contractile phenotype where actomyosin-derived contractile forces maintain vascular tone (7). However, VSMCs display remarkable plasticity and during vessel remodelling, they undergo phenotypic transition to a synthetic, migratory phenotype. This synthetic phenotype is prevalent in vascular disease, including atherosclerosis and aneurysm (10, 11). Phenotypic modulation is associated with changes in VSMC contractile protein expression and morphology (12); however, we still do not fully understand whether the ability of VSMCs to generate actomyosin derived-force is altered by VSMC phenotypic modulation.

Microenvironment rigidity transmits ‘outside-in’ forces to VSMCs and this process is dependent on adhesions that convey force between the ECM and cytoskeleton (13–16). VSMCs and other cell types respond to outside-in signals by exerting actomyosin based contractile forces on the matrix (inside-out forces) that scale with ECM stiffness (13, 15, 16). Rho/ROCK signalling is rapidly activated at ECM adhesions in response to matrix rigidity, to augment actomyosin activity, via actin polymerisation and myosin light chain phosphorylation (15). Actin cytoskeletal reorganisation and enhanced actomyosin activity increase VSMC integrated traction forces, the force VSMCs apply to the ECM. In other cell types, matrix rigidity promotes enhanced adhesion and actomyosin activity to increase traction force (17–20). However, our knowledge of the balance between actomyosin activity, adhesion organisation and traction force magnitude in response to matrix rigidity in VSMCs remains limited. Traction force microscopy (TFM) remains the gold standard for measuring traction forces generated by cells, but is highly specialised and time consuming to perform. As such, only a limited number of studies have measured VSMC traction force. Those that have been performed have reported that matrix rigidity promotes increased traction force magnitude and adhesion reorganisation (21–23). However, the mechanisms driving these changes remain unknown. Therefore, in this current study we investigate the impact of matrix rigidity on VSMC traction force generation. We show that VSMC spreading and integrated traction force (integrated-TF) magnitude display a moderate correlation and that this relationship was independent of matrix rigidity. Importantly, we show that actomyosin activity and adhesion number are inversely regulated by matrix rigidity and VSMC spreading. This inverse regulation results in a net zero change in TF/μm^2^. However, VSMC spreading results in increased adhesive contractile units per cell which augments integrated-TF. Our study suggests that VSMC traction force is tightly regulated by a compensatory mechanism involving differential actomyosin and adhesion regulation.

## Methods

### Cell culture

Human adult aortic VSMCs (passage 3-10) were purchased from Cell Applications Inc (Isolate-1). Human adult aortic 35F VSMCs were a kind gift from Professor Cathy Shanahan (King’s College London, UK) (Isolate-2). VSMCs were grown in growth media (Cell Applications Inc) and were otherwise cultured as described previously (24).

### Polyacrylamide hydrogel preparation and Traction Force Microscopy

Hydrogels were prepared as described previously (25, 26). A JPK Nanowizard-3 atomic force microscope was used to confirm hydrogel stiffness as described previously (25). VSMCs were seeded onto 12kPa hydrogels containing 0.5μm red fluorescent (580/605) FluoSpheres (Invitrogen). Imaging was performed using a Nikon Eclipse Ti-E live cell imaging system to capture 20x magnification images before and after cell lysis by the addition of 0.5% Tx-100. Drift was corrected using the ImageJ StackReg plugin and traction force was calculated using the ImageJ plugin described previously to measure FluoSphere displacement (27). Briefly, bead displacement was measured using the first and last image of the movie sequence. To determine integrated (cell total) the cell region was segmented by overlaying the traction map with the cell image, highlighting the cell traction region with an ROI and extracting the traction forces in each pixel by using the save XY coordinate function in ImageJ. To generate gradient trend line data, integrated-TF was plotted against VSMC area on a scatter plot in Graph Pad Prism 7. Linear regression analysis was performed to generate:

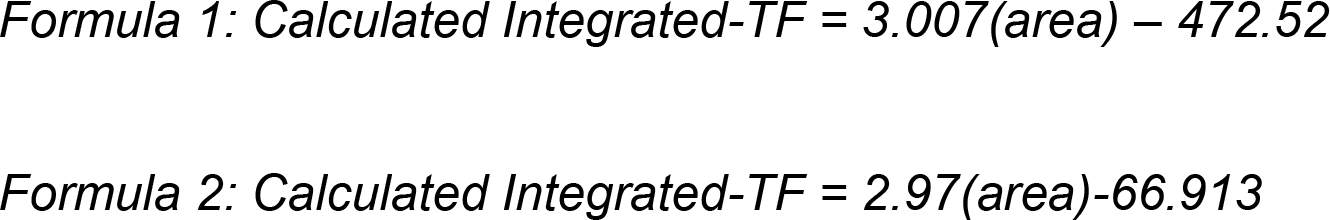

### Western blot analysis

Antibodies used for WB, IF; Vinculin (V9131), α-actin, β-actin (A5316) (Sigma). Secondary antibodies for WB were horseradish peroxidase-conjugated anti mouse (NA931) or anti rabbit (NA94V) antibodies from GE Healthcare. ECL chemiluminescent kit (RPN2132, GE Healthcare) was used for detection according to manufacturer’s instructions.

### Fixed cell microscopy and data analysis

Cells were cultured on hydrogels, fixed in paraformaldehyde and processed as described previously (28). Invitrogen anti-mouse Alexa fluor 568 (A11031) and anti-rabbit Alexa fluor 488 (A11034) were used as IF secondary antibodies. Rhodamine Phalloidin was used to stain F-actin and DAPI was used to visualise cell nuclei. All images were captured at 20x magnification using a Leica SP2 laser scanning confocal microscope. Cell area and aspect ratio was measured using the ImageJ open source analysis software (28). pMLC mean intensity and total intensity were measured using the ROI function in ImageJ. Focal adhesion number and size were measured in ImageJ, as described previously (28).

### Statistics

Results are presented as mean +/− SEM. Results are presented to show data distribution and each point corresponds to any individual measurement. For comparison of matrix stiffness group’s, paired student t-tests were performed. GraphPad Prism 7 software was used to generate trend-line gradient equations, scatter plots and to perform linear regression and statistical analysis.

## Results

### Matrix stiffness stimulates VSMC spreading and increases traction force magnitude

We set out to validate previous findings showing that matrix rigidity stimulates spreading and actomyosin activity in VSMCs. VSMCs were grown on collagen-1 coated polyacrylamide hydrogels, with an average Young's modulus of 12 (healthy) and 72kPa (stiff) (Supplementary Figure 1A). Western blot analysis revealed that levels of smooth muscle actin (SM-actin), calponin and β-actin remained unchanged by matrix stiffness (Supplementary Figure 1B), suggesting that matrix rigidity does not alter expression of VSMC contractile proteins. Immunofluorescence microscopy (IF) analysis confirmed that VSMCs were larger on 72kPa hydrogels compared to those grown on 12kPa hydrogels (Supplementary Figures 1C and D). Aspect ratio remained unchanged by matrix stiffness (Figures 1C and E). To observe cellular force generation, we performed traction force microscopy on VSMCs grown on 12 and 72kPa hydrogels. The ability of VSMCs to displace beads embedded in the hydrogels was reduced on 72kPa compared to 12kPa hydrogels (Figures 1A and B). However, integrated VSMC traction force (integrated-TF) magnitude was enhanced on the 72Kpa hydrogels compared to 12kPa hydrogels (Figures 1A and C). These data confirm previous findings showing that matrix rigidity influences VSMC spreading and traction force.

**Figure 1.**
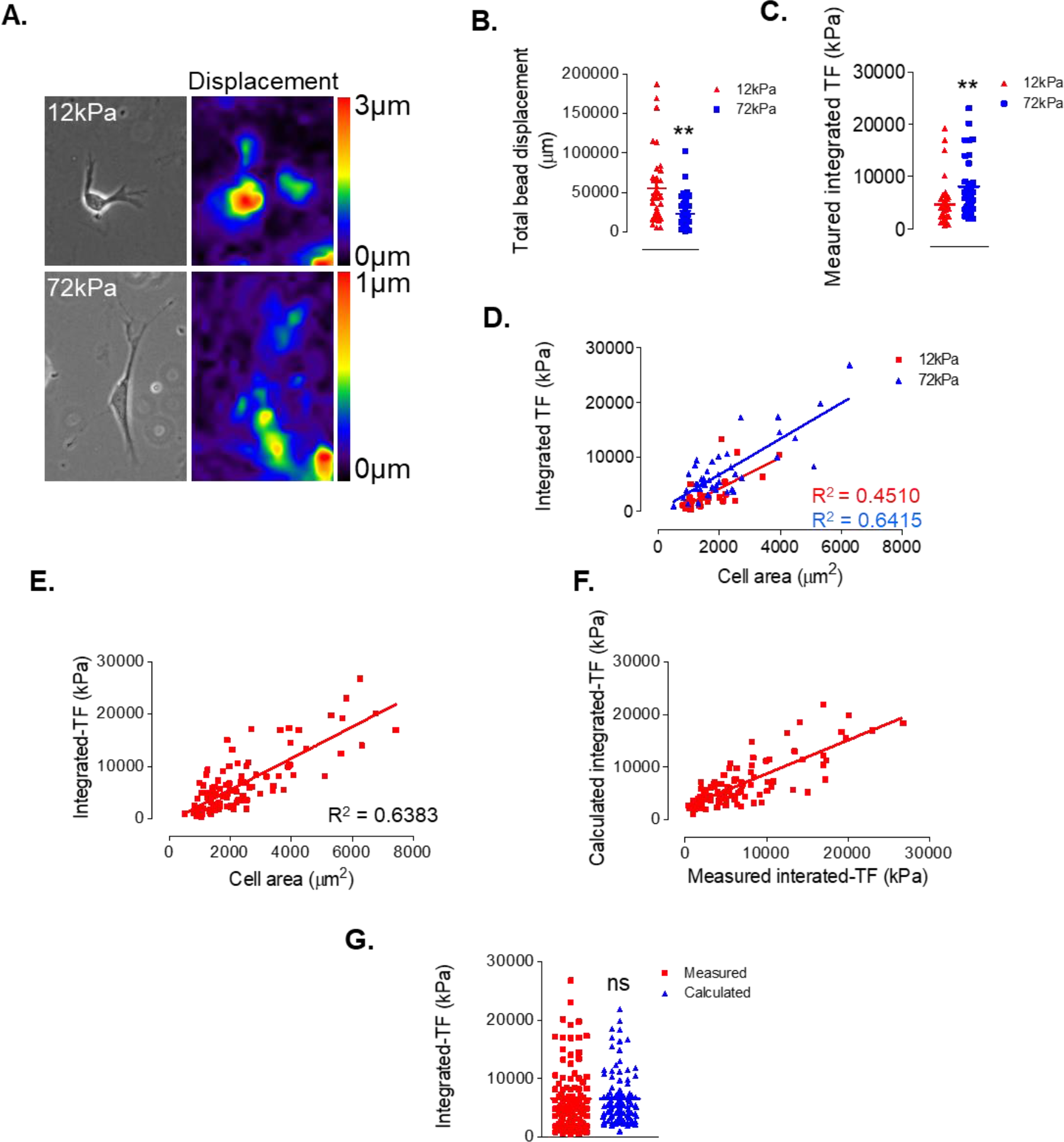
VSMC spreading/integrated-TF magnitude relationship is matrix rigidity independent. **A)** Representative bead displacement maps for isolate-1 VSMCs grown on 12 and 72kPa hydrogels. Graphs show **B)** integrated bead displacements and **C)** integrated-TF magnitude for VSMCs grown on 12 (n=29) and 72kPa (n=42) hydrogels. **D)** Graph shows distribution of integrated-TF magnitude plotted against VSMC area on 12kPa (red) (n=29) and 72kPa (blue) (n=42) hydrogels. **E)** Graph shows combined distribution of integrated-TF plotted against VSMC area on 12kPa and 72kPa hydrogels (n=104). Graphs show **F**) distribution of calculated versus measured integrated-TF magnitude and **G)** comparison of measured and calculated integrated-TF using the trend-Line formula (formula 1) generated in **E**). (** p = <0.01 and ns = non-significant).

### VSMC spreading predicts integrated-TF magnitude independently of matrix rigidity

The above data shows that matrix rigidity enhances both VSMC spreading and integrated traction force magnitude. VSMC spreading has previously been reported to correlate with traction force magnitude on soft 1kPa hydrogels (29). We next speculated that matrix rigidity influences the relationship between VSMC spreading and traction force magnitude. Analysis revealed that a moderate correlation between VSMC spreading and integrated-TF magnitude existed on both 12kPa (R^2^ = 0.4510) and 72kPa (R^2^ = 0.6415) hydrogels (Figure 1D). Importantly, the relationship between VSMC spreading and integrated-TF was not significantly altered by matrix rigidity, suggesting that changes in traction force magnitude were driven by changes in VSMC spreading.

As the VSMC spreading/integrated-TF magnitude relationship was independent of matrix rigidity, we hypothesised that integrated-TF could be calculated from VSMC area. To test this idea, we combined the data from the 12kPa and 72kPa hydrogels and calculated the gradient of the trend line (Figure 1E). The formula generated was used to calculate integrated-TF magnitude from the measured VSMC area. As expected, the spread of calculated data closely resembled that of measured integrated-TF magnitude (Figures 1F and G). To test this hypothesis further, we used an independent aortic VSMC isolate. TFM was performed to experimentally measure VSMC traction force. The measured data confirmed that bead displacement was reduced whereas integrated-TF magnitude was enhanced by matrix rigidity, recapitulating the above findings (Figures 2A-C). Similar to our above findings, the integrated-TF/spreading relationship for isolate-2 displayed no significant difference between 12kPa (R^2^ = 0.4818) and 72kPa (R^2^ = 0.6325) (Figure 2D). The estimated integrated-TF was independently calculated from VSMC area, using the trend line formula generated from isolate-1. Comparison of the measured and calculated integrated-TF revealed no significant difference between the two data sets (Figure 2E). Next, we sought to refine our model and compared the measured integrated-TF/spreading relationship between the two independent isolates. Both isolates displayed a similar integrated-TF/spreading relationship (Figure 2F), and as there was no significant difference between the two isolates, we combined all the data and calculated the gradient of the trend line (Figure 2G). This combined formula was used to calculate the integrated-TF of both isolates. Comparison revealed that the measured and calculated integrated-TF, were in close agreement with no significant difference between these data sets (Figures 2H and I).

**Figure 2.**
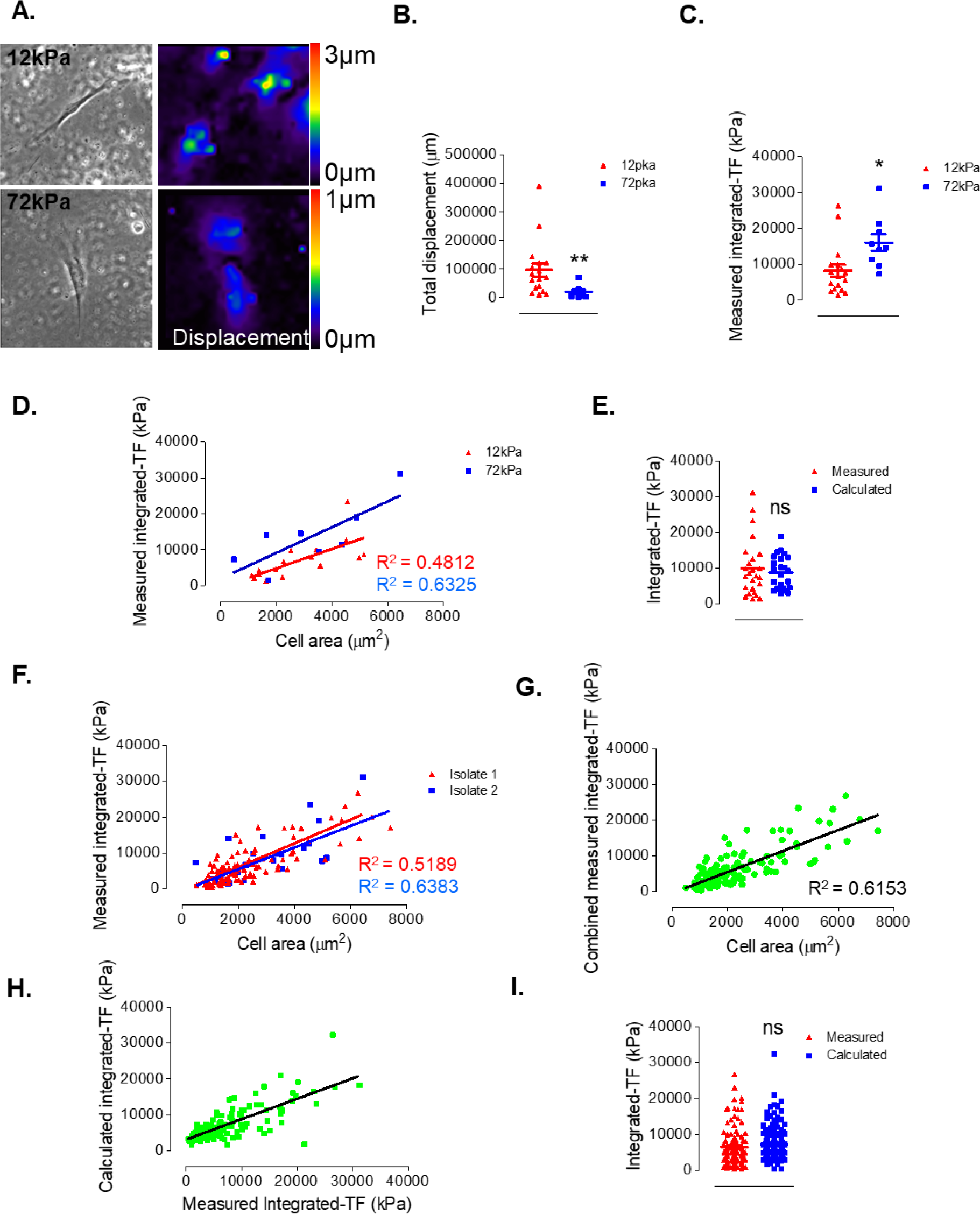
Refinement of the estimated integrated-TF formula. **A)** Representativebead displacement maps for isolate-2 VSMCs grown on 12 and 72kPa hydrogels. Graphs show **B)** integrated bead displacements and **C)** integrated-TF magnitude for isolate-2 grown on 12 (n=17) and 72kPa (n=8) hydrogels. **D)** Graph shows distribution of measured integrated-TF magnitude plotted against VSMC area for isolate-2 grown on 12kPa (red) (n=17) and 72kPa (blue) (n=9) hydrogels. **E)** Graph shows distributionof measured and calculated integrated-TF of isolate-2 using formula 1. **F)** Graph shows distribution of measured integrated-TF magnitude versus VSMC area of isolate-1 (red) (n=104) and isolate-2 (blue) (n=25). **G)** Graph shows the combined distribution of measured integrated-TF and area of isolate-1 and isolate-2 used to generate formula 2 (n=129). **H)** Graph shows the combined distribution of calculated versus measured integrated-TF for isolate-1 and isolate-2. **I**) Graph shows the distribution of measuredand calculated integrated-TF magnitude for isolate-1 and isolate-2 combined data. (*p = <0.05, **p = <0.01 and ns = non-significant).

### MLC phosphorylation strongly correlates with calculated integrated-TF magnitude

To further confirm that VSMC spreading and calculated integrated-TF correlated, we used an independent marker of actomyosin activity. We next performed immunofluorescence microscopy to assay phosphorylated myosin light chain (pMLC) levels in VSMCs grown on 12kPa and 72kPa hydrogels. Surprisingly, analysis revealed that pMLC mean intensity was reduced on 72kPa compared to 12kPa hydrogels, however, the total pMLC intensity remained unchanged (Figures 3A-C). Further analysis confirmed that VSMC spreading and the calculated integrated-TF magnitude strongly correlated with the total pMLC intensity on both 12kPa (R^2^ = 0.8658) and 72kPa (R^2^ = 0.9384) hydrogels (Supplementary Figure 2 and Figure 3D). However, there was no correlation between pMLC mean intensity and VSMC area/estimated integrated-TF (Supplementary Figure 2 Figure 3E). The R^2^ values were not significantly different between 12kPa and 72kPa hydrogels for the relationship between total pMLC total intensity and VSMC spreading/calculated integrated-TF magnitude. This suggests that total pMLC levels was also independent of matrix stiffness.

**Figure 3.**
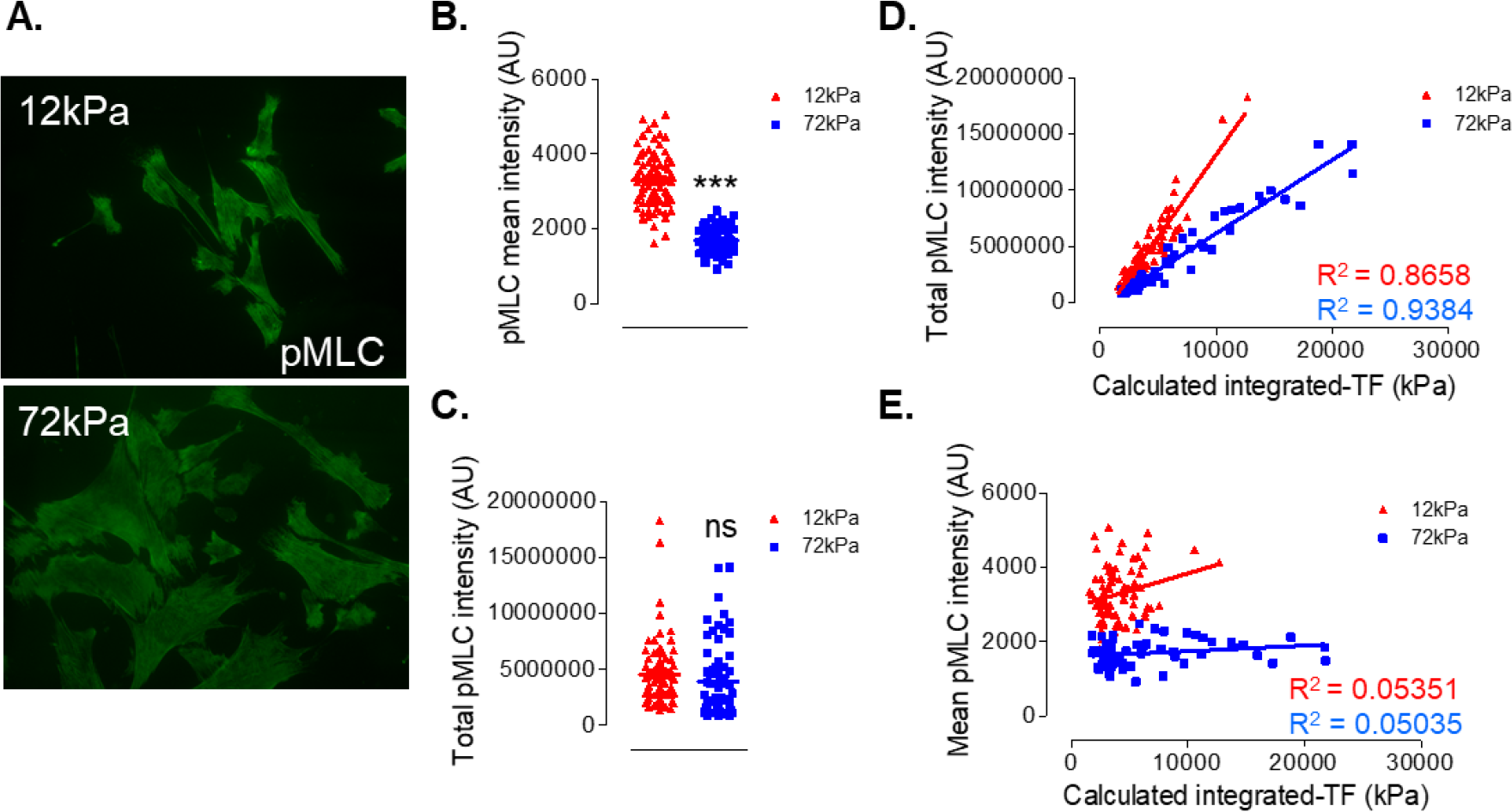
Total pMLC levels correlate with calculated integrated-TF. A) Representative images of pMLC stained isolate-1 VSMCs grown on 12 and 72kPa hydrogels. Graphs show B) pMLC mean intensity and C) pMLC total intensity in isolate-1 VSMCs grown on 12 (n=77) and 72kPa (n=61) hydrogels. Graphs show D) pMLC mean intensity and E) pMLC total intensity verses calculated integrated-TF on 12 (red) and 72kPa (blue) hydrogels. (***p = <0.0001 and ns = non-significant).

### Increased cell: matrix adhesion number enhances VSMC traction force production

The above data shows that suggests that increased VSMC spreading was not stimulating enhanced actomyosin-generated forces. We predicted a role for VSMC matrix adhesions, so we performed IF analysis of VSMCs stained with an antibody raised to the matrix-adhesion protein vinculin. Analysis revealed that VSMCs grown on 72kPa hydrogels possessed an increased number of matrix adhesions and larger matrix adhesions compared to VSMCs grown on 12kPa hydrogels (Figure 4A-C). This data shows that VSMC matrix adhesions are altered by matrix rigidity. Next, using formula 2, the integrated-TF was calculated from the cell area data and confirmed that cell area and calculated integrated-TF were enhanced by matrix rigidity (Supplementary Figure 3 and Figure 4D). We next examined the relationship between VSMC spreading/calculated integrated-TF magnitude and vinculin organisation. Analysis revealed that VSMC spreading/calculated integrated-TF magnitude correlated with the number of vinculin adhesions but not adhesion size (Figures 4E, F and Supplementary Figure 3). Analysis revealed a significant difference between the 12kPa and 72kPa trend-line gradient, confirming that this relationship is enhanced by matrix rigidity (Figures 4E, F and Supplementary Figure 3). Finally, we sought to dissect whether increased integrated-TF magnitude was driven by increased adhesion number in larger VSMCs. Analysis revealed that the average-TF per adhesion was decreased on 72kPa hydrogels (Figure 4G). Importantly, VSMCs grown on 72kPa hydrogels possessed an increased number of adhesions/μm^2^ (Figure 4H) and therefore displayed a similar TF/μm^2^ as VSMCs on 12kPa hydrogels (Figure 4I).

**Figure 4.**
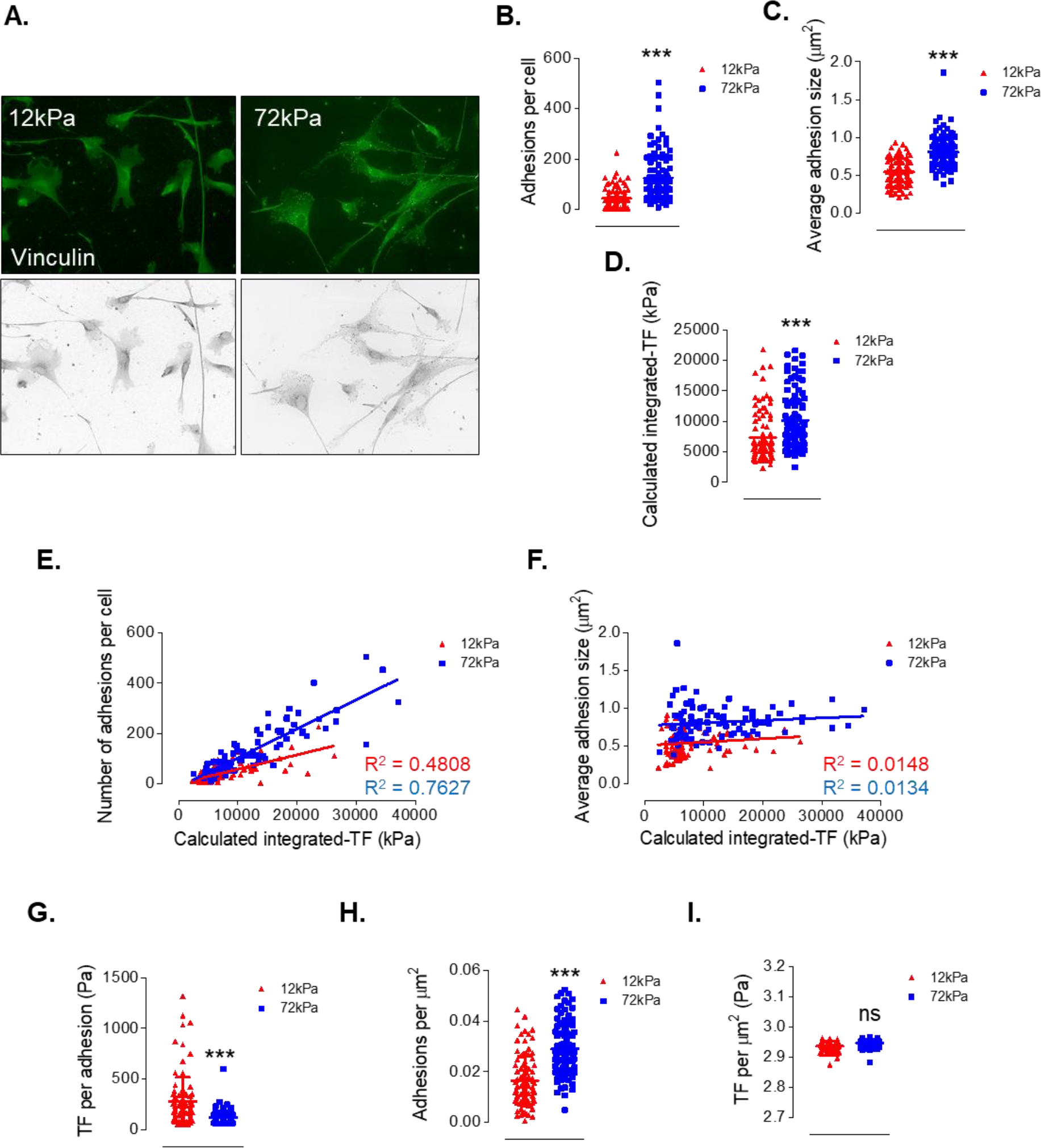
Matrix rigidity influences VSMC matrix adhesion organisation. A) Representative images of vinculin stained isolate-1 VSMCs grown on 12 and 72kPa hydrogels. Graphs show B) average adhesion size and C) number of adhesions per cell on 12kPa (n=94) and 72kPa (n=99) hydrogels. Graphs represent the combined data from 3-independent experiments. (***p= <0.0001). Graphs show D) adhesion size and E) adhesion number per cell verses calculated integrated-TF on 12 (red) and 72kPa (blue) hydrogels. Graphs show F) calculated integrated-TF, G) integrated-TF per adhesion, H) adhesion number/μm2 and I) integrated-TF/me for VSMCs grown on 12 and 72kPa hydrogels. (***p = <0.0001 and ns = non-significant).

## Discussion

Decreased aortic compliance is observed during vascular ageing and disease, however, our understanding of how changes in aortic stiffness influence VSMC function remains limited. Decreased aortic compliance is ultimately driven by changes in the ECM composition, including elastin fragmentation and collagen accumulation (5, 6). VSMCs normally adopt a quiescent, contractile phenotype in the vessel wall, however, VSMCs are not terminally differentiated and can adopt a synthetic phenotype (30). Matrix rigidity has been shown to promote VSMC proliferation, suggesting that microenvironment rigidity influences VSMC phenotype (30). Matrix rigidity is also known to influence VSMC morphology and traction force magnitude however, the mechanisms driving these processes remain to be defined (31, 32). In agreement with previous findings using immature embryonic aortic VSMCs, we show that VSMC traction force magnitude and spreading are enhanced by matrix rigidity. However, this previous study used hydrogels between 10kPa and 25kPa (32). These rigidities are within the normal range of adult aortic rigidity (4). Our study is the first to examine this phenomenon in mature adult VSMCs at a rigidity more akin to disease. In other cell types, matrix rigidity enhances the relationship between traction force magnitude and cell spreading (33). However, we show that this relationship in adult VSMCs is independent of matrix rigidity in two independent VSMC isolates. Furthermore, we have generated a model for estimating integrated-TF magnitude from VSMC spreading data. TFM is a time consuming and specialised technique that can only be performed on live cells. We have used our model to estimate integrated-TF magnitude using fixed VSMC data. Importantly, we show that in a population of smooth muscle cells there is no significant difference in between measured and calculated integrated-TF.

Although increased VSMC spreading stimulated integrated-TF magnitude, TF/μm^2^ remained unaltered by both matrix rigidity and spreading. This suggests that actomyosin-generated force that is transferred to the ECM by VSMCs is unaffected by spreading. However, pMLC analysis revealed that matrix rigidity reduces pMLC levels, suggesting that actomyosin activity is reduced by matrix rigidity. Matrix adhesions remodel in response to matrix rigidity and changes in both adhesion number per cell and adhesion size have been reported in a variety of cell types (34). VSMC matrix adhesion number increased with spreading, yet the average-TF per adhesion was reduced. Importantly, matrix rigidity stimulated increased matrix adhesion numbers which resulted in equivalent levels of TF/μm^2^, as VSMCs grown on 12kPa hydrogels. These data suggest that enhanced VSMC anchor points and not enhanced actomyosin activity contribute to the increase in integrated-TF. Our data also shows that matrix rigidity inversely regulates pMLC levels and matrix adhesion number in VSMCs; matrix rigidity promotes decreased pMLC levels and increased adhesion anchor points. Matrix stiffness also increases VSMC spreading, however, due to this inverse pMLC/adhesion number regulation, there is no change on force transfer/μm^2^ between VSMCs and the ECM. As a result, the integrated-TF/VSMC spreading relationship is matrix rigidity independent. This suggests that the force/μm^2^ exerted by VSMCs on the ECM is tightly regulated. We propose that matrix rigidity triggers a compensatory mechanism to preserve force/μm^2^ that potentially protects VSMCs from actomyosin-induced damage (Figure 5). Unrestrained actomyosin activity induces DNA damage in other cells types and DNA damage drives VSMC ageing (35–37). Therefore, this compensatory mechanism potentially protects VSMC from actomyosin-induced ageing.

**Figure 5.**
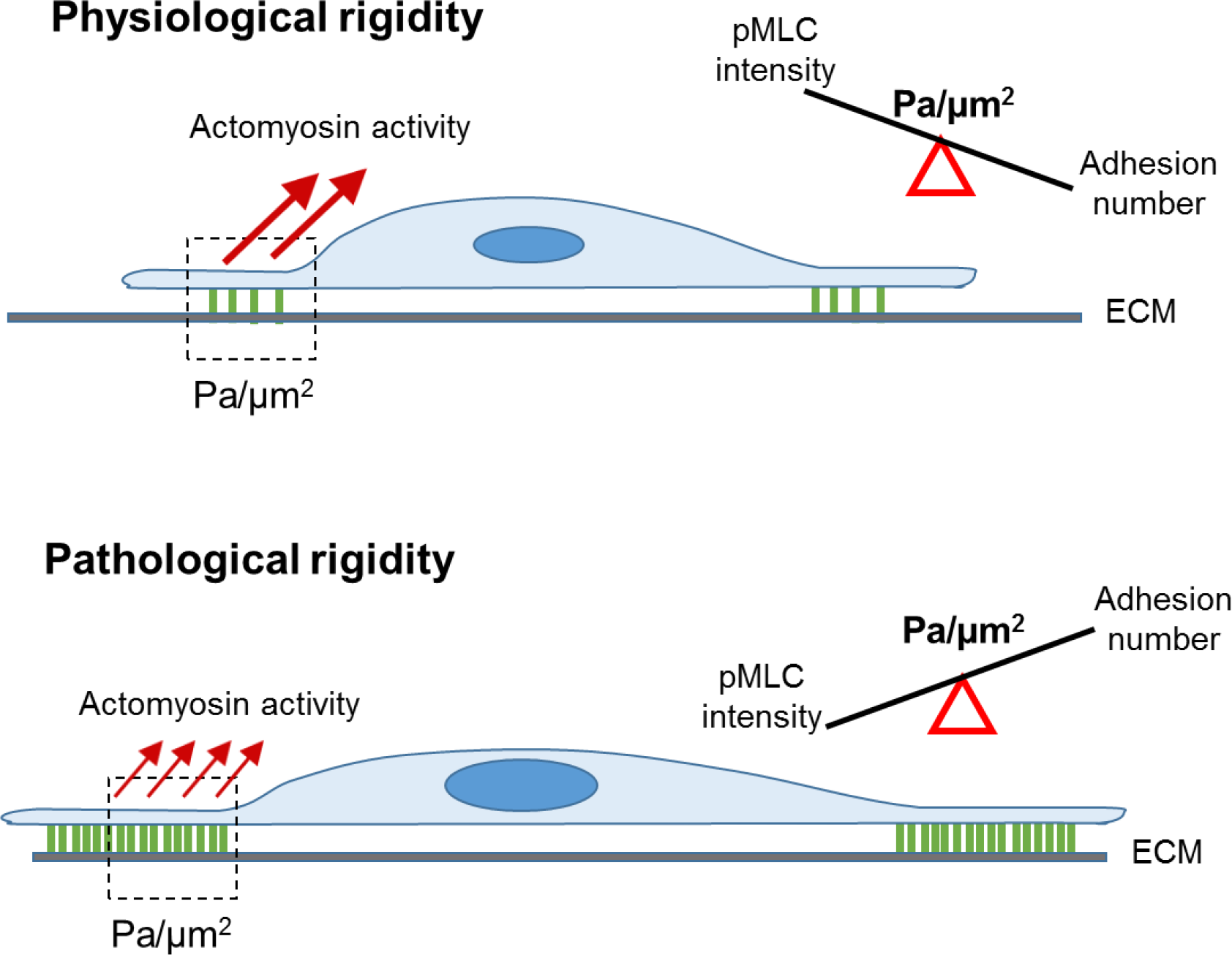
Model representing how differential adhesion number and actomyosin regulation preserve VSMC integrated-TF/μm^2^.

Despite this compensatory mechanism, VSMC spreading promotes increased adhesive contractile units that ultimately increase integrated-TF. However, despite generating increased integrated-TF, the ability of VSMCs to deform the ECM diminishes with matrix rigidity. Our findings confirm that VSMCs can efficiently displace the ECM at normal (12kPa) aortic stiffness and this is in agreement with the role of VSMCs in the maintenance of vascular tone (7). Enhanced actomyosin activity deforms the surrounding ECM and enables VSMCs contraction resulting in decreased vascular compliance. However, our findings show that a 6-fold increase in stiffness dramatically diminishes the ability of VSMCs to deform their microenvironment. These findings suggest that despite matrix rigidity stimulating VSMC actomyosin activity, a 6-fold increase in aortic stiffness would result in the VSMCs being unable to significantly contribute to vascular tone and compliance. Changes in VSMC phenotype are known to influence expression of smooth muscle cell contractile proteins (38). For example, contractile VSMCs possess both smooth muscle (SM)-myosin II and non-muscle (NM)-myosin II, whereas synthetic VSMCs possess predominantly NM-myosin II (39). Although structurally similar, SM-myosin II and NM-myosin II possess different force generating capabilities/functions; NM-myosin II generates less force than SM-myosin II and NM-myosin II contributes to tonic contraction, whereas SM-myosin contribute to phasic contraction (40–42). Our understanding of the relative contribution of NM-myosin II and SM-myosin II in VSMC dysfunction remains unknown. Synthetic VSMCs were used in this study and we cannot rule out the possibility that contractile VSMCs possess a greater ability to deform stiffened matrix than their synthetic counterparts. However, given that matrix stiffness has been demonstrated to promote VSMC proliferation, the relationship between VSMC force generation, phenotype and vessel compliance is likely to be complex. As aortic stiffness and VSMC phenotype switching are observed in vascular diseases, including atherosclerosis and hypertension, further research is now needed to better understand the relationship between VSMC force generation and vessel compliance. Dramatic changes in cytoskeletal organisation and morphology are associated with VSMC phenotypic transition and our model will allow further interrogation of these pathways that influence VSMC force generation during disease associated phenotypic modulation.

## Funding

This work was supported by a British Heart Foundation (BHF) Non-clinical PhD Studentship (FS/17/32/32916).

## Author Contributions

SA and PM conducted experiments and performed data analysis. DTW designed the experiments and wrote the manuscript.

## Conflict of Interest

Conflict of interest: none declared.

## Data availability

The raw/processed data required to reproduce these findings cannot be shared at this time due to technical or time limitations.

