## Supplementary Figures for "A model for estimating traction force magnitude reveals differential regulation of actomyosin activity and matrix adhesion number in response to smooth muscle cell spreading"

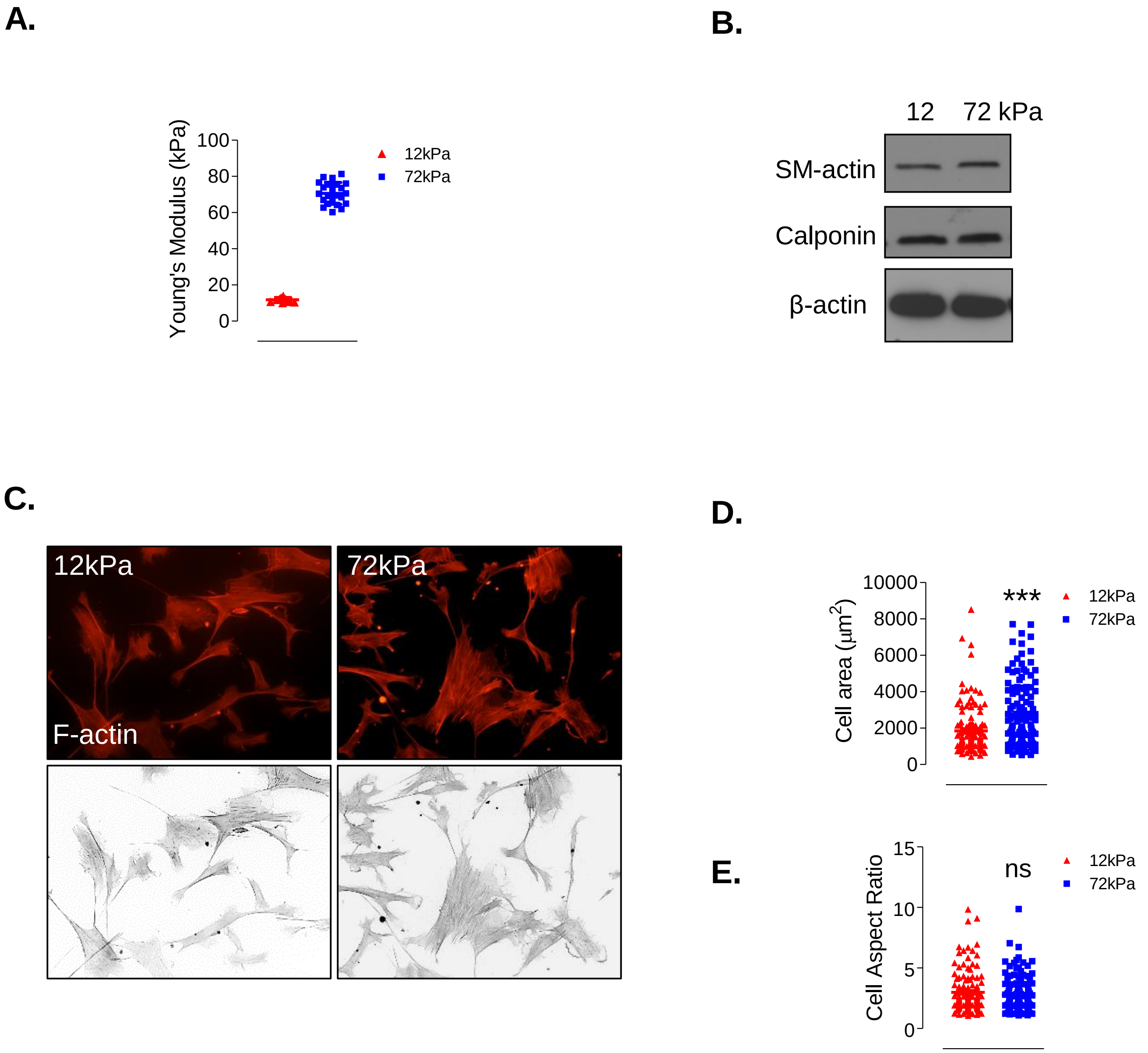


**Supplementary Figure 1.** Matrix rigidity influences VSMC spreading. **A)** Characterisation of the Young's modulus of hydrogels used in this study. **B)** WB of smooth muscle cell marker expression in VSMCs grown on 12 and 72kPa hydrogels. **C)** Representative images showing F-actin staining of VSMCs grown on 12 and 72kPa hydrogels. Graphs show **D)** cell area and **E)** cell aspect ratio. Graphs represent the combined data of 3-independent experiments analysing >200 VSMCs (***p= <0.0001 and ns = non-significant).

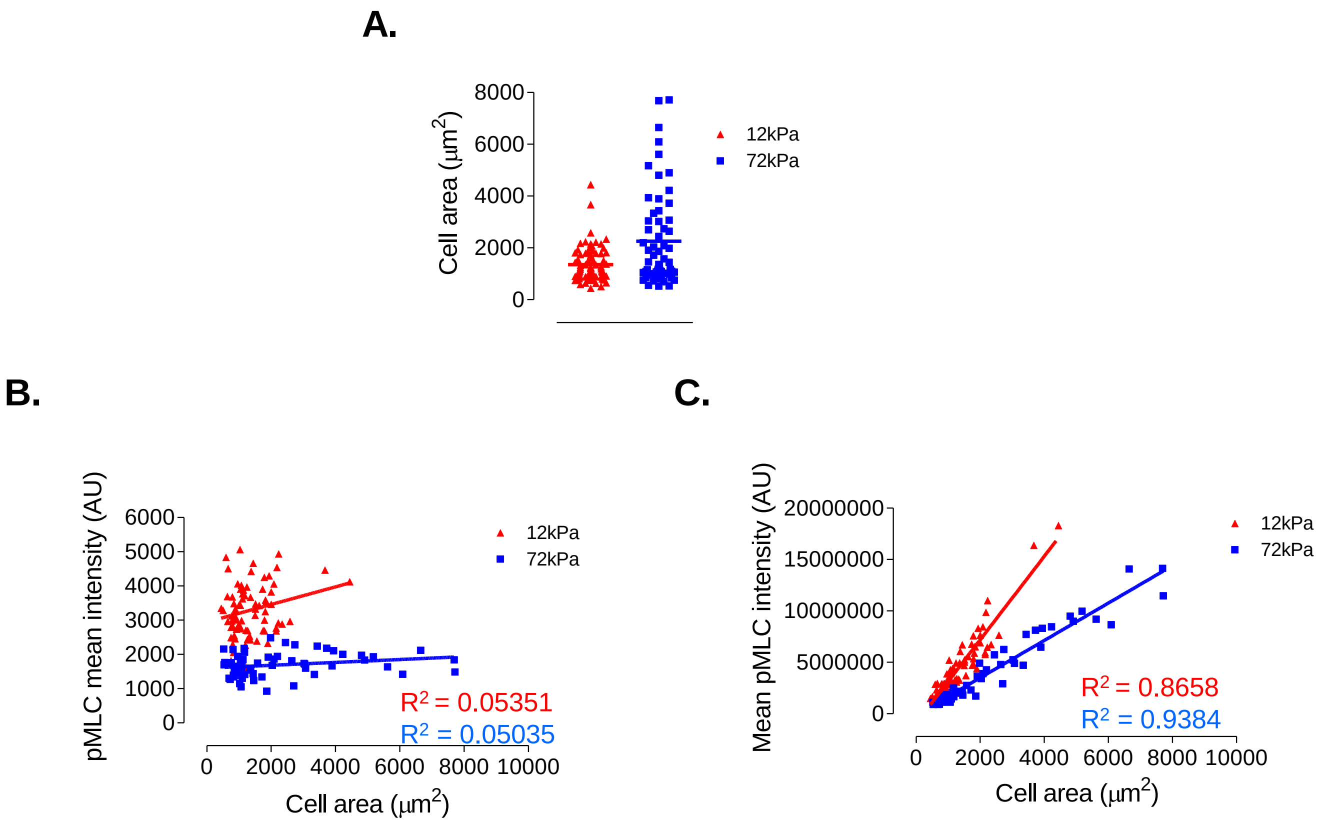


**Supplementary Figure 2.** Total pMLC levels correlate with VSMC spreading. **A)** Graph shows cell area of pMLC stained isolate-1 VSMCs grown on 12 and 72kPa hydrogels. Graphs show **B)** pMLC mean intensity and **C)** pMLC total intensity in isolate-1 VSMCs grown on 12 and 72kPa hydrogels.

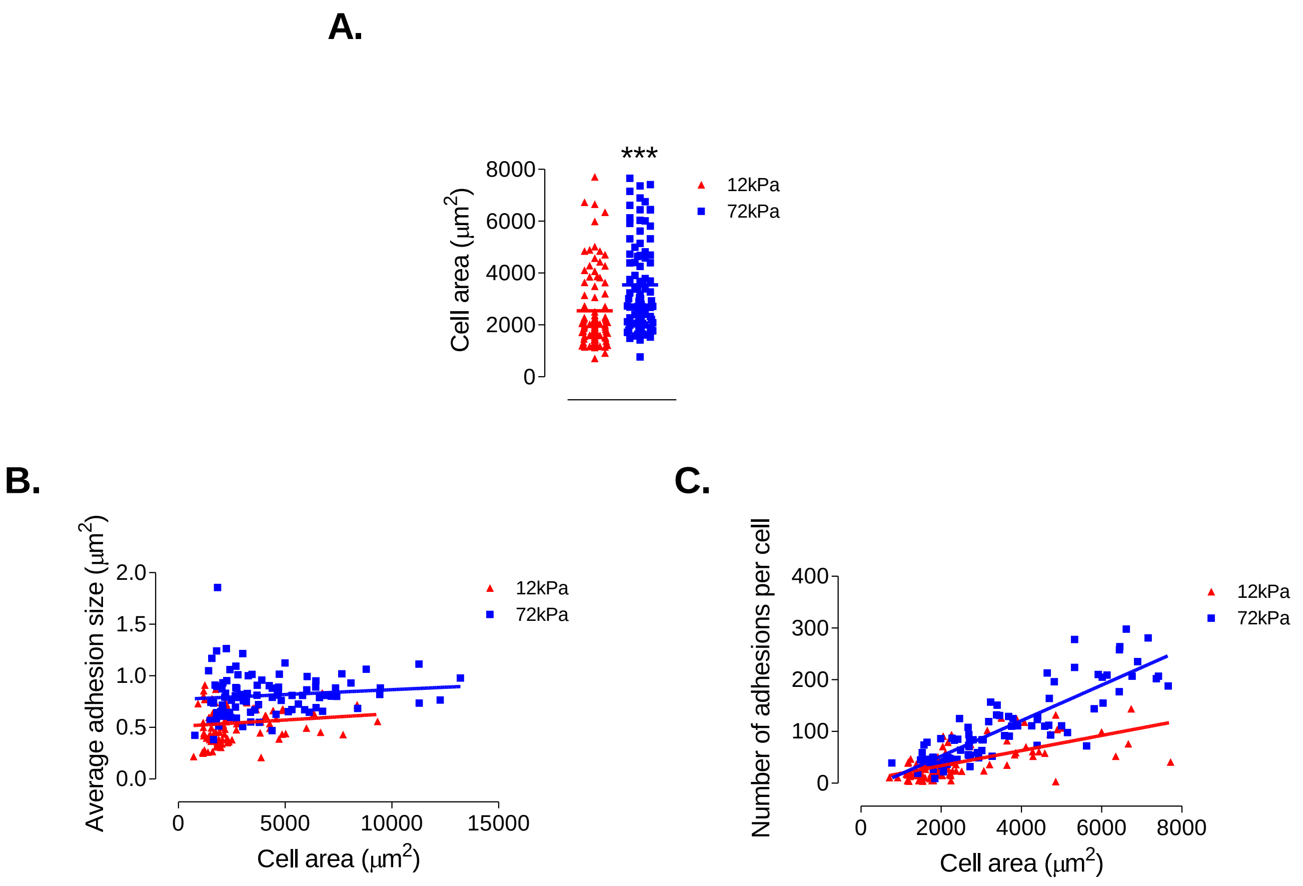


**Supplementary Figure 3.** Adhesion number correlates with VSMC spreading. **A)** Graph shows cell area of vinculin stained isolate-1 VSMCs grown on 12 and 72kPa hydrogels. Graphs show **B)** adhesion size and **C)** adhesion number per cell verses isolate-1 VSMC area on 12 (red) and 72kPa (blue) hydrogels.
